# Mechanistic modeling suggests stroma-targeting antibody-drug conjugates as an alternative to cancer-targeting in cases of heterogeneous target expression

**DOI:** 10.1101/2025.01.31.635847

**Authors:** N Ezgi Wood, Anil Cengiz, Ming Gao, Alexander V Ratushny, Ronny Straube

**Affiliations:** Quantitative Pharmacology, Data, and Analytics, Bristol Myers Squibb, Princeton, NJ, USA; Department of Mathematics, University of Utah, Salt Lake City, UT, USA; Quantitative Pharmacology, Data, and Analytics, Bristol Myers Squibb, Cambridge, MA, USA; Quantitative Pharmacology, Data, and Analytics, Bristol Myers Squibb, Seattle, Washington, USA; Pioneering Medicines, Cambridge, MA, USA

## Abstract

Antibody-drug conjugates (ADCs) are gaining increasing traction in the treatment of oncological diseases; however, many clinical failures have also been observed. One key factor limiting ADC effectiveness is the heterogeneous expression of their target antigen. While the vast majority of ADCs in clinical development target antigens on cancer cells (cancer-targeting), they can also target antigens expressed on non-cancerous stromal cells in the tumor microenvironment (stroma-targeting). It remains unclear whether targeting cancer cells or stromal cells is more effective in suppressing tumor growth. Here, we present three related mathematical models to evaluate: (1) cancer-targeting ADCs with homogeneous target antigen expression, (2) cancer-targeting ADCs with heterogeneous target antigen expression, and (3) stroma-targeting ADCs. Our simulations suggest that cancer-targeting ADCs can achieve high efficacy when their target antigen is homogenously expressed. However, in cases of heterogeneous antigen expression, cancer-targeting ADCs may lead to an initial reduction in tumor size, followed by regrowth due to the elimination of antigen-positive cells and expansion of antigen-negative cells. This limitation could potentially be overcome by stroma-targeting ADCs, as antigen-positive stromal cells may continue to be recruited into the tumor by the oncogenic factors produced by the remaining cancer cells. Furthermore, we demonstrate that ADCs with more permeable payloads and less stable linkers may offer improved efficacy in the context of heterogeneous target expression.

## Introduction

Antibody-drug conjugates (ADCs) are therapeutic agents composed of three main components: an antibody, a payload, and a linker that connects the two. This combination enables ADCs to leverage the specificity of antibodies for targeting and potency of the payloads to kill the target cells (Tsuchikama et al., 2024). Upon binding to their targets on the cell surface, ADCs are internalized, and the payload is released inside the cells. The payload can then diffuse out of the target cells into neighboring cells and kill them, a phenomenon known as bystander killing. This mechanism allows ADCs to eliminate antigen-negative (Ag-) cells in the tumor microenvironment (TME) as well (Figure 1A) (Dumontet et al., 2023).

**Figure 1:**
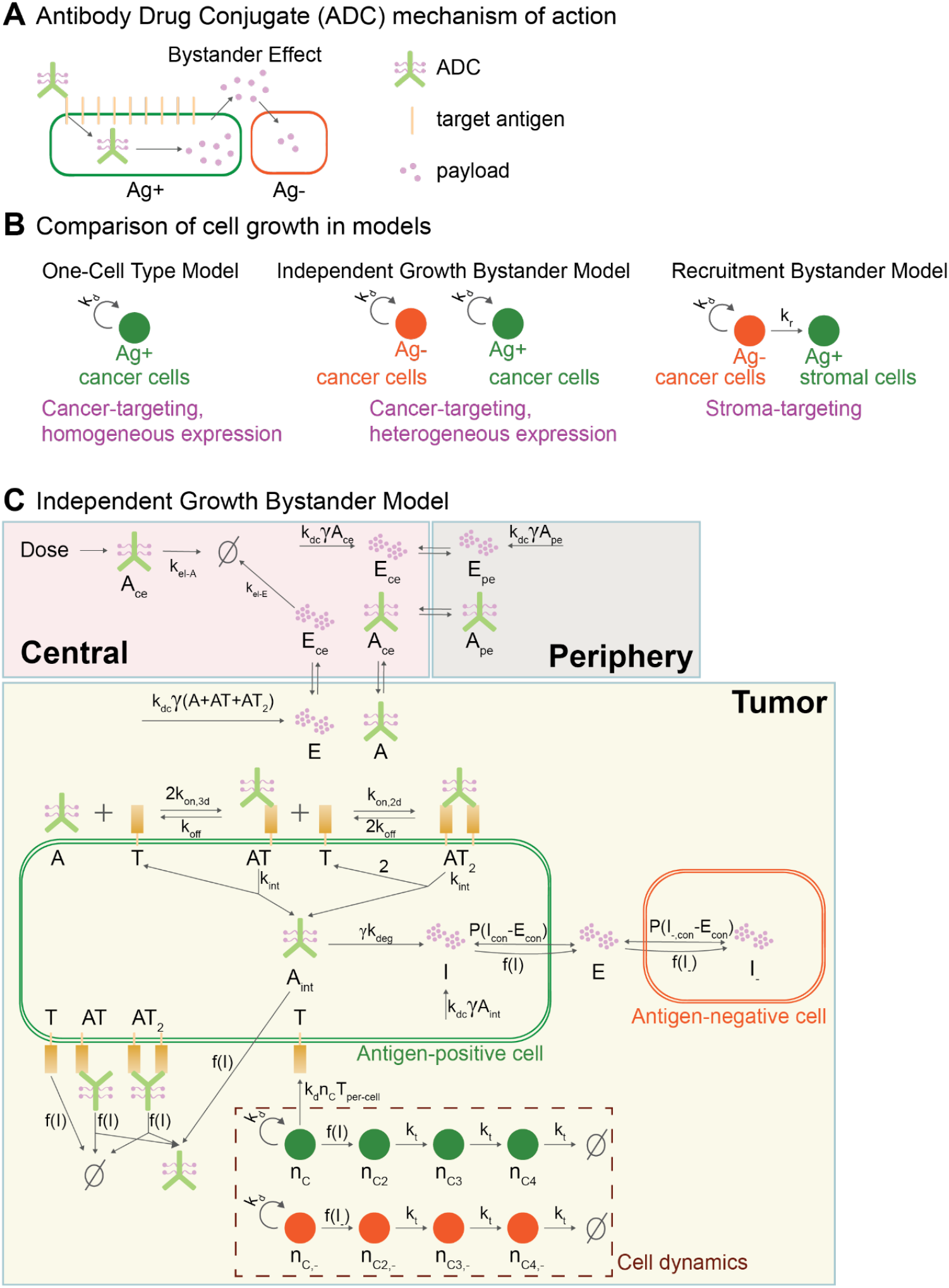
Overview of the bystander effect and the models. **A)** ADC mechanism of action and bystander effect. **B)** Cell growth dynamics in models: 1) One-Cell Type Model: represents a cancer-targeting ADC directed at a tumor with homogeneous target expression among cancer cells. 2) Independent Growth Bystander Model: represents a cancer-targeting ADC aimed at a tumor with heterogeneous target expression among cancer cells. 3) Recruitment Bystander Model: represents a stroma-targeting ADC, where the target is only expressed by stromal cells and not by cancer cells. **C)** Overview of the Independent Growth Bystander Model. Subscript con denotes concentration. Functions f(I) and f(I_-_) denote sigmoidal functions of the intracellular payload per cell (see Supplementary). Note that the One Cell Model does not include Ag-cells (Supplementary Figure 1). The Recruitment Bystander Model differs from the Independent Growth Bystander Model only in cell growth dynamics, where the growth of Ag+ cells depends on the Ag-cells (Supplementary Figure 2).

ADCs can target antigens expressed on cancer cells (cancer-targeting ADC) or on non-cancerous cells within the TME, such as stromal cells (stroma-targeting ADC) (Diamantis & Banerji, 2016; Fu et al., 2022; Xiao & Yu, 2021). Cancer-targeting ADCs with heterogeneous expression of their target antigen, as well as stroma-targeting ADCs, rely on bystander effects to kill Ag-cancer cells. Since antigen expression heterogeneity is one of the significant factors leading to poor treatment outcomes (Grairi & Le Borgne, 2024; Tsuchikama et al., 2024), optimizing ADCs for bystander efficacy is crucial for addressing this challenge. However, it remains unclear whether cancer-targeting or stroma-targeting ADCs are more effective in bystander killing, or how ADCs can be optimized to enhance bystander killing.

Predicting ADC clinical outcomes, optimizing their properties, and selecting the appropriate therapeutic indications are challenging tasks due to the complex and often counterintuitive behavior of ADCs (Colombo et al., 2024; Colombo & Rich, 2022; Dumontet et al., 2023). Mechanistic mathematical models have proven instrumental in unraveling the complex relationships between ADC properties and ADC efficacy, thereby guiding the development of optimal ADC compounds for clinical application (Lam et al., 2022).

To address these challenges and evaluate bystander effects under various scenarios, here we present three related mechanistic models for ADCs. These models enable comparisons between cancer-targeting ADCs with homogeneous or heterogeneous target antigen expression and stroma-targeting ADCs.

Building on existing mechanistic models for ADC modality (Byun & Jung, 2019; Khera et al., 2018; Scheuher et al., 2022; Shah et al., 2012; Singh & Shah, 2019), we expand the framework to encompass both Ag+ and Ag-cells, as well as key ADC mechanisms of action, including binding to target antigen, crosslinking on the cell surface, payload release, payload diffusion across the cell membrane, and linker deconjugation. Our models distinguish between cancer-targeting and stroma-targeting ADCs based on interactions between Ag+ and Ag− cells. For cancer-targeting ADCs with heterogeneous target expression, we assume Ag+ and Ag-cancer cell populations grow independently. In contrast, for stroma-targeting ADCs, we model Ag+ stromal cell growth as dependent on the Ag-cancer cells, given that stromal cells are present in the TME due to the oncogenic drivers (Feng et al., 2022; Zhang et al., 2023) and that a disrupted TME will be reconstructed by tumor cells (Xiao & Yu, 2021).

Our simulations indicate that cancer-targeting ADCs may have limited long-term efficacy if their target is expressed heterogeneously. Although the tumor might initially shrink, it will eventually regrow due to expansion of Ag-cancer cells. This limitation arises because ADC treatment leads to a higher intracellular payload concentration in Ag+ cells, rendering them more sensitive to ADC treatment. However, ADCs targeting stromal cells might overcome this limitation, as Ag+ stromal cells may be recruited to the tumor by oncogenic drivers produced by the remaining Ag-cells.

Additionally, our results highlight that optimal ADC properties depend on target antigen expression patterns. For homogeneous target expression on cancer cells, cancer-targeting ADCs with less permeable payloads may achieve better efficacy. Conversely, for heterogeneous target expression, more permeable payloads may be more effective. Lastly, we demonstrate that a certain degree of linker instability can enhance ADC efficacy, suggesting the existence of an optimal level of linker stability for maximizing therapeutic outcomes.

## Methods

### Overview of mathematical models

Here, we present three related ordinary differential equation (ODE) models describing the mechanism of action of ADCs in various scenarios. The first model is the One-Cell Type Model, which incorporates only Ag+ cells in the Tumor compartment, representing Ag+ cancer cells. This model simulates a cancer-targeting ADC aimed at a tumor with homogenous target expression in the cancer cells (Figure 1B, Supplementary Figure 1). The second and third models are bystander models that incorporate both Ag+ and Ag-cells in the Tumor compartment. In the Independent Growth Bystander Model, both Ag+ and Ag-cells represent cancer cells that grow independently, simulating a cancer-targeting ADC applied at a tumor with heterogeneous target expression (Figure 1B, C). In contrast, in the Recruitment Bystander Model, Ag+ cells represent the stromal cells expressing the target, while Ag-cells represent the cancer cells lacking the target (Figure 1B, Supplementary Figure 2). In this model, Ag+ cell growth is driven by Ag-cancer cells to incorporate the relation between cancer and stromal cells. This model simulates a stroma-targeting ADC where the target is only expressed by stromal cells.

In all models, the pharmacokinetics of the ADC and the payload are captured using a two compartment PK framework that includes Central and Peripheral compartments with elimination occurring in the Central compartment (Figure 1C). To accurately represent the PK of the ADC and the payload, different Central and Peripheral volumes are used for each, following a similar approach to that described in (Shah et al., 2012).

The models incorporate the diffusion of ADC and payload between the Central and Tumor compartments. Within the Tumor compartment, the models account for ADC binding to its target, ADC crosslinking on the cell surface (avidity), ADC internalization, ADC degradation and payload release. ADC degradation and payload release in the cells are modeled as a single step. The payload can diffuse in and out of the cells, and in the bystander models, it can also diffuse in and out of Ag-cells. In the models, the number of targets per cell is constant and the total number of targets is linearly proportional to Ag+ cell number. Cell dynamics are modeled using the exponential-linear growth and the damaged cell compartments introduced by (Simeoni et al., 2004). Tumor size is linked to the total cell number.

In the Recruitment Bystander Model, Ag+ cell growth depends on the presence of Ag-cells. The initial ratio of Ag-to Ag+ cells is assumed to reflect the steady-state ratio that these populations reach prior to treatment. This ratio is then used to set the rate of recruitment of Ag+ cells by Ag-cells (Supplementary Section 2).

Spontaneous linker deconjugation is also included and is assumed to occur at the same rate in every compartment. In the models, the ADC represents the total antibody of any drug-to-antibody ratio (DAR), while the change in the average DAR is modeled separately to account for the linker deconjugation process. We show that if the linker deconjugation rate is uniform across all conjugation sites, the average DAR decreases exponentially (Supplementary Section 5).

### Model parametrization

Model parameters and equations are given in Supplementary Material. The models are parametrized using literature values based on Trastuzumab Deruxtecan (Supplementary Table 4). For payload PK parameters we use exatecan PK values (Garrison et al., 2003). The parameters were selected within a physiological range to facilitate a meaningful exploration of the models’ key properties.

### Model implementation

The models are implemented in SimBiology and simulated using MATLAB 2022a. The model sbproj file is included in the Supplementary Material.

## Results

### Cancer-targeting ADCs may provide limited durability of response when the target antigen is expressed heterogeneously, whereas stroma-targeting ADCs may overcome this limitation

Here, we simulate 5.4 mg/kg Q3W ADC treatment schedule. This treatment regimen leads to tumor regression with the One-Cell Type Model (Figure 2A).

**Figure 2:**
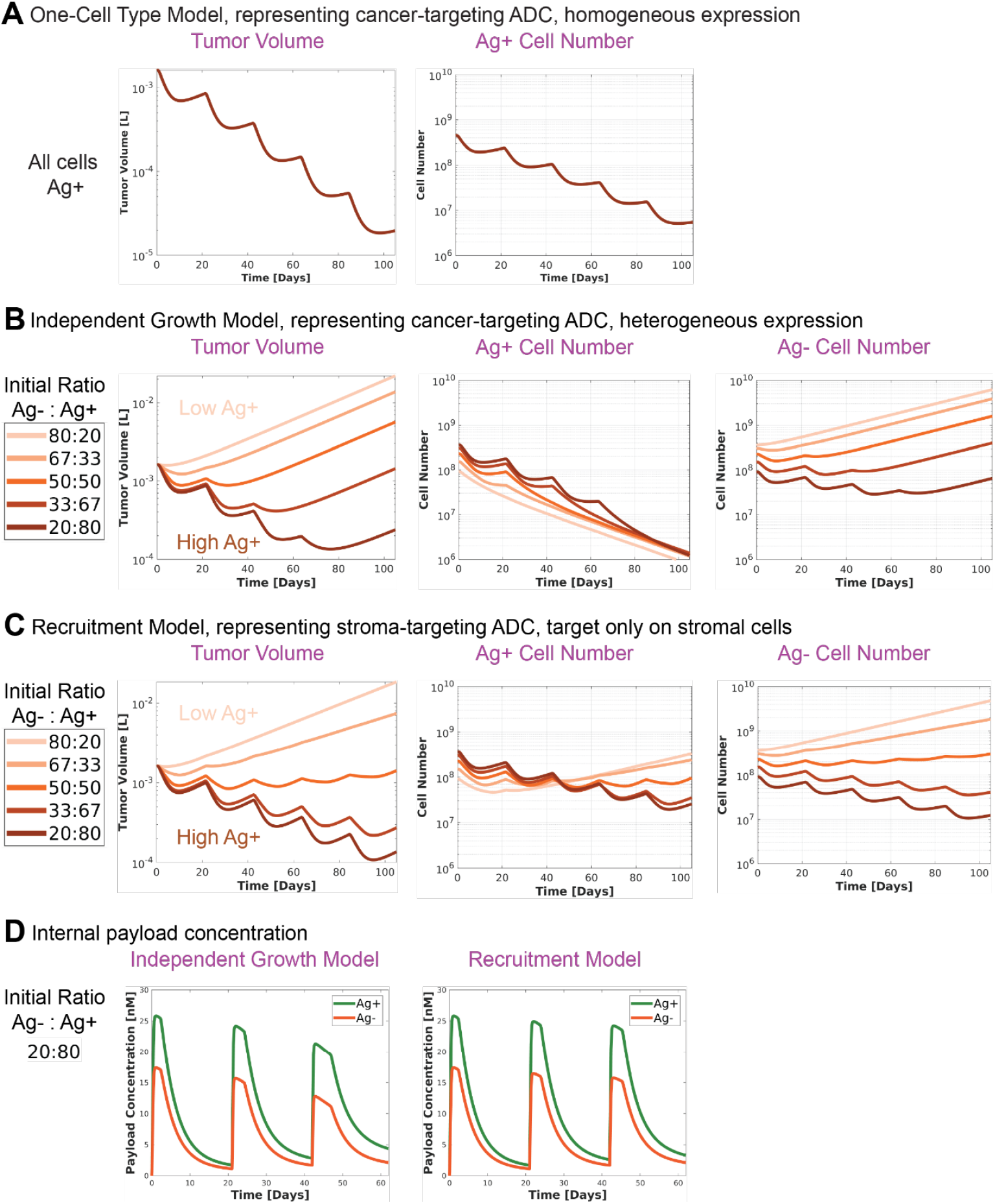
Comparison of cell growth and intracellular payload concentrations in bystander models. **A)** One-Cell Type Model. Tumor shrinks. **B)** Independent Growth Bystander Model. Tumor eventually grows across all tested initial Ag-:Ag+ ratios. **C)** Recruitment Bystander Model. Durable tumor suppression can be achieved when the initial percentage of Ag+ cells is sufficiently high. **D)** Intracellular payload concentrations are higher in Ag+ cells compared to Ag-cells.

In the Independent Growth Bystander Model, which represents cancer-targeting ADCs with heterogeneous target expression, the treatment does not achieve a durable response for all initial Ag-to Ag+ ratios with the current model parameters. Eventually, the tumor regrows (Figure 2B). A higher initial percentage of Ag+ cells delays regrowth. While the number of Ag+ cells consistently decreases, the tumor regrowth is driven by the proliferation of Ag-cells. In contrast, in the Recruitment Bystander Model, representing a stroma-targeting ADC, durable tumor regression is achievable for sufficiently high initial percentage of Ag+ cells (Figure 2C).

The internal payload dynamics elucidate the limitations of ADC mechanism of action in the bystander setting: ADCs initially enter the Ag+ cells and the payload is released in these cells. The payload then diffuses out of Ag+ cells into the TME and subsequently into Ag-cells, down its concentration gradient. This diffusion process results in a higher internal payload concentration in Ag+ cells compared to Ag-cells, making Ag+ cells more sensitive to ADC treatment (Figure 2D).

In deterministic ODE models, cell numbers are treated as continuous variables, thus, the Ag-cell population in the Independent Growth Bystander Model always recovers and grows back. However, in reality, the Ag-cell population may become extinct. To simulate this scenario, a threshold can be introduced, assuming the Ag-cell population becomes extinct if it falls below this threshold. For illustration, we have chosen a threshold of 10^7 cells, which is below the detection limit of current methods (Crosby et al., 2022; Frangioni, 2008). Under the current model parametrization, an initial Ag+ cell population of approximately 90% is required to drive Ag-cell population below this threshold within 84 days (Supplementary Figure 3).

### Mouse xenograft models support model predictions

Admixed xenograft models, where a mixture of Ag+ and Ag-cells are injected into mice, provide an experimental validation for the simulations with the Independent Growth Bystander Model (Figure 2B). The findings from admixed xenograft models reported by (Li et al., 2016) align with the model projections: tumors containing a mixture of Ag+ and Ag-cells eventually regrow. As the percentage of Ag+ cells increases, the regrowth is delayed further (Figure 7 in (Li et al., 2016)). The authors also report that after treatment, only Ag-cells remain (Figure 5 in (Li et al., 2016)).

Xenograft models with Ag-cancer cells and stroma-targeting ADCs serve as a test case for the simulations with the Recruitment Bystander Model (Figure 2C). In the study by (Purcell et al., 2018), an Ag-cancer cell line was treated with a stroma-targeting ADC, allowed to grow, and then retreated.

Unlike the admixed xenograft models, the tumor regrowth in this case included Ag+ cells. Additionally, retreating the tumor resulted in a similar degree of tumor regression as the initial treatment (Figure 4C in (Purcell et al., 2018)). This mirrors the simulation results (Figure 2C), where Ag+ cells are not totally depleted with the Recruitment Bystander Model, as long as Ag-cells remain present.

### Less permeable payloads may enhance efficacy in tumors with homogeneous target expression, whereas more permeable payloads may be more effective in tumors with heterogeneous target expression

Next, we investigate the optimal properties of ADCs for maximum efficacy in homogeneous vs heterogeneous settings. First, we vary the permeability of the payload across the cell membrane. In the One-Cell Type Model, where every cell is Ag+, increasing payload permeability results in decreased efficacy (Figure 3A). In contrast, in the bystander models, increasing payload permeability leads to increased efficacy (Figure 3B, C).

**Figure 3:**
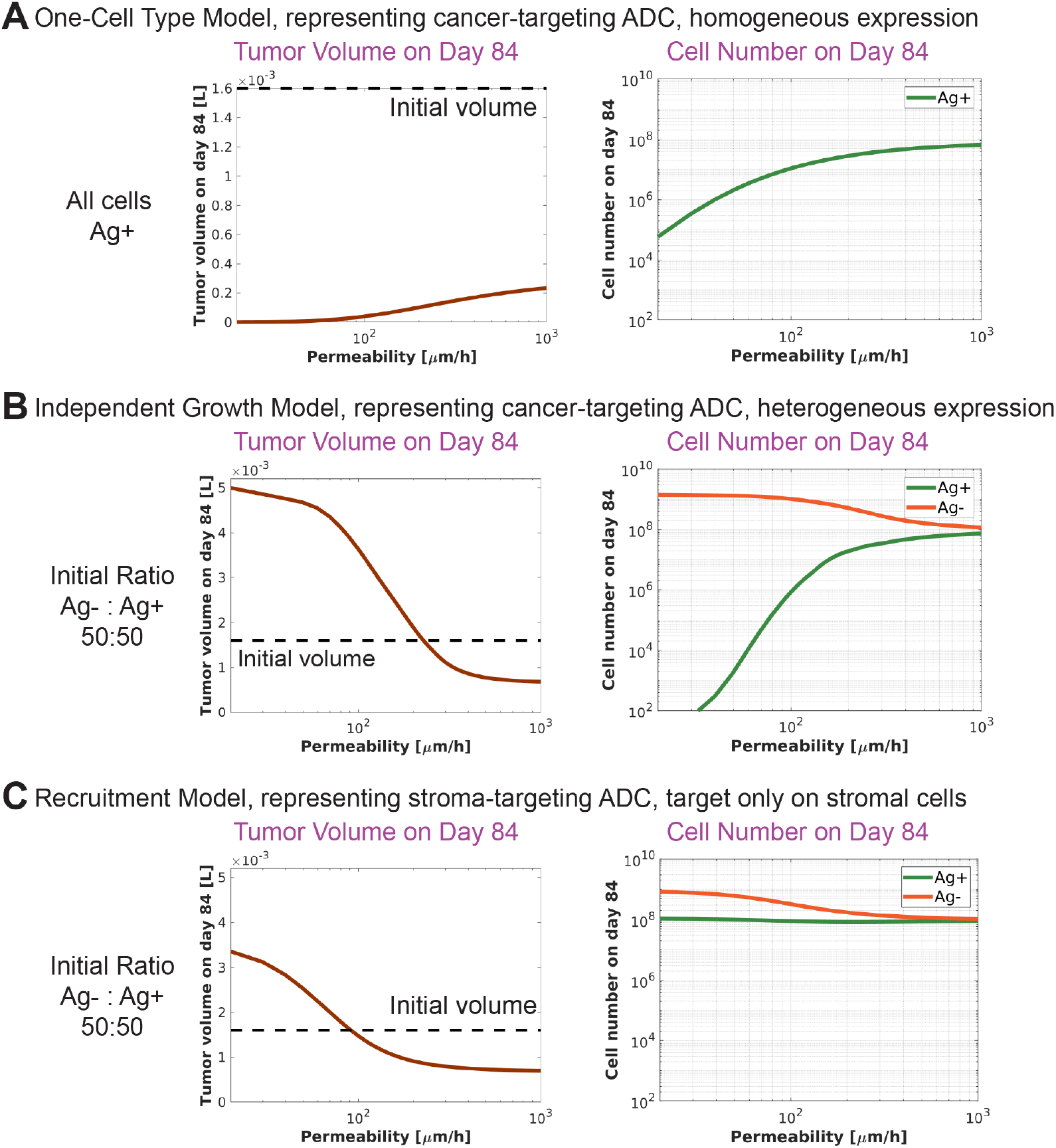
Effect of payload membrane permeability on ADC efficacy. **A)** One-Cell Type Model. As payload permeability increases, ADC efficacy decreases. **B)** Independent Growth Bystander Model. **C)** Recruitment Bystander Model. In both **B)** and **C)**, payload membrane permeability is varied for both Ag+ and Ag-cells. For bystander models, as payload permeability increases, the ADC efficacy increases. Note that payload permeability may have an optimal value for a different set of parameters or time points (see Supplementary Figure 5).

**Figure 4:**
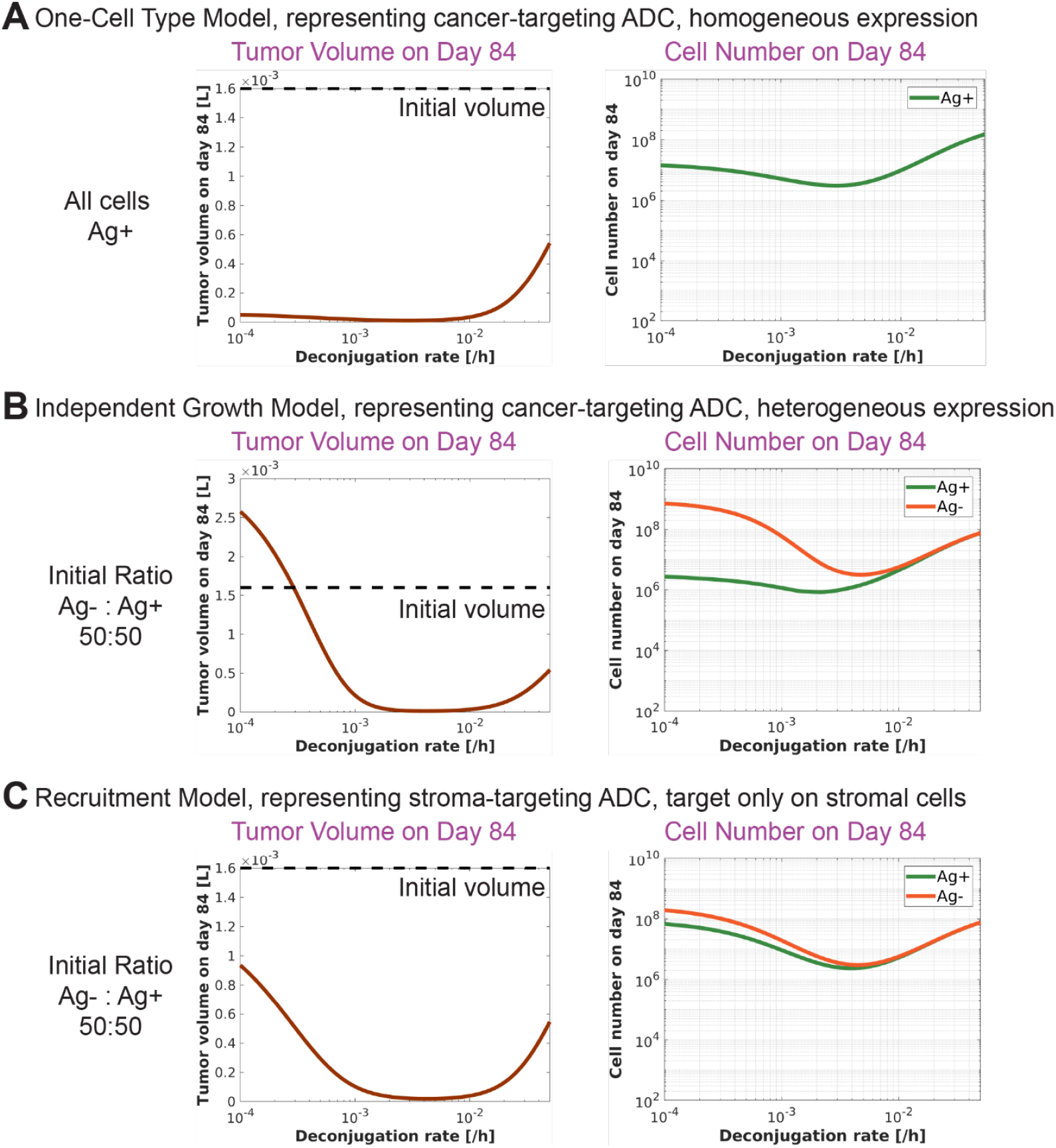
Effect of linker stability on ADC efficacy. **A)** One-Cell Type Model. **B)** Independent Growth Bystander Model. **C)** Recruitment Bystander Model. In all models, a decrease in linker stability, i.e. an increase in deconjugation rate, initially enhances efficacy but eventually reduces it. This effect is more pronounced in the bystander models.

Increased payload permeability allows for more payload to diffuse out of Ag+ cells, resulting in less efficient killing of these cells. Conversely, increased payload permeability facilitates more efficient transport of the payload into the Ag-cells, leading to more efficient killing of Ag-cells. Thus, in the bystander setting, increasing permeability might lead to increased Ag+ cell number while decreasing the number of Ag-cells (Figure 3B). As a result, the Ag+ population might be maintained for a longer period, and Ag-cells can be killed more effectively, leading to better tumor suppression. Therefore, the permeability of the payload across cell membrane could be critical in determining whether a tumor shrinks or grows (Figure 3B, C).

Note that in the bystander models, for different parameter regions or other time points, the opposing effect of payload permeability on the killing of Ag+ and Ag-cells may result in an optimal payload permeability value for maximum efficacy (Supplementary Figure 4).

### Linker deconjugation may increase ADC efficacy

Payload release through spontaneous linker deconjugation can either increase or decrease ADC efficacy. Increasing the rate of spontaneous linker deconjugation may elevate the levels of free payload in the TME, which could lead to higher internal payload concentrations within the cells by reducing the payload concentration gradient between TME and the cells. However, spontaneous linker deconjugation also reduces the average DAR, resulting in less payload being delivered to the cells. Additionally, the release of the payload from ADC can lead to faster elimination of the payload from the system, as payload clearance occurs faster than ADC clearance.

Here, we simulate various linker deconjugation rates to assess their impact on efficacy. Across all models, there is an optimal linker deconjugation rate that maximizes efficacy by balancing the aforementioned effects (Figure 4). The impact of linker deconjugation on efficacy is more pronounced in the bystander models that the One-Cell Type model (Figure 4A vs B,C).

While a less stable linker may improve efficacy, it can also increase toxicity due to the elevated levels of free payload in circulation (Nguyen et al., 2023). Therefore, linker stability should be optimized taking both efficacy and toxicity into consideration.

## Discussion

In this study, we present novel models to assess ADC efficacy across various scenarios, providing key insights into ADC mechanism of action and potential limitations of this modality. Our simulations indicate that cancer-targeting ADCs may not provide durable responses when their target antigen is heterogeneously expressed. Specifically, ADCs cause higher internal payload levels in Ag+ cells than Ag-cells. While this leads to initial tumor regression, the subsequent reduction in Ag+ cell population allows the Ag-cells to drive tumor regrowth. Stroma-targeting ADCs are a promising alternative to overcome this limitation, under the assumption that oncogenic drivers produced by the remaining Ag-cells will recruit the Ag+ stromal cells to the tumor.

In our simulations with bystander models, the same payload sensitivity and permeability parameters were used for the Ag+ and Ag-cells to keep the focus on the dynamics generated by cell interactions, avoiding confounding factors introduced by parameter differences. Similarly, in the Independent Growth Bystander Model, identical growth parameters were used for both cell types. In practice, however, these parameters could be different for Ag+ and Ag-cells. For example, in an admixed xenograft mouse model, Ag+ cells overexpressing the target might grow more slowly, while knockout Ag-cells might grow faster, ultimately reducing ADC efficacy.

Furthermore, using the same payload sensitivity parameters for both Ag+ and Ag-cells might underestimate the efficacy of stroma-targeting ADCs, as stromal cells might be less sensitive to cytotoxic payloads like microtubule inhibitors or Topoisomerase 1 inhibitors, which predominantly affect rapidly dividing cells (Jordan & Wilson, 2004; Tomicic & Kaina, 2013). Thus, the efficacy of the stroma-targeting ADCs might be better than projected if accounting for such differences. Another factor affecting stroma-targeting ADC efficacy is the Ag+ stromal cell to Ag-cancer cell ratio. While stromal cells may be more abundant than cancer cells in certain tumor types, as reflected in tumor-stromal ratio (TSR) (van Pelt et al., 2018), the proportion of target bearing stromal cells is likely smaller than the overall TSR. A lower percentage of Ag+ stromal cells would reduce the effectiveness of stroma-targeting ADCs. These potential parameter differences between Ag+ and Ag-cells underscore the importance of calibrating the models to the specific ADC and patient population under investigation.

In the models presented here the payload moves down its concentration gradient. Potential effects of efflux pumps, which could alter the relative payload concentrations in Ag+ and Ag-cells, are not considered. Additionally, for the Recruitment Bystander Model, we sought to include the most basic relationship that would recapitulate existing literature data, thus, more complex relationships between cancer cells and stromal cells are omitted.

Target antigen expression on cancer cells often exist along a continuum. Here, we consider two scenarios: one where all cancer cells express the target and another with two distinct subpopulations — one expressing the target and the other lacking it. However, there could be multiple subpopulations of cancer cells exhibiting varying levels of target expression, ranging from low to high. The models presented here represent the two limiting cases. Additionally, some targets may be expressed on both cancer cells and stromal cells (Purcell et al., 2018), which would enhance the efficacy predicted by our models. While our current focus is on these limiting cases, the models can be readily extended to more complex scenarios.

Targeting stomal cells may also trigger immune-related mechanisms. For instance, the elimination of stromal cells could enhance immune cell infiltration and boost efficacy (Xiao & Yu, 2021). However, this study primarily examines bystander killing and does not address immune system-related effects.

Additionally, while the ODE models assume a well-mixed cell population, the spatial distribution of cells could be non-uniform, and spatial effects may influence efficacy. These spatial effects are beyond the scope of this study.

Our simulations indicate that optimal ADC properties depend on the target expression pattern. While a less permeable payload might be more effective for a homogeneously expressed target, a more permeable payload could achieve better efficacy with a heterogeneously expressed target. Similarly, a less stable linker may improve efficacy. While our models focus on efficacy, clinical candidate optimization requires a careful balance between efficacy and toxicity. Thus, an unstable linker, though beneficial for efficacy, might increase toxicological risks, rendering it unsuitable for clinical development. Overall, our results underscore the promise of stroma-targeting ADCs as a viable alternative to cancer-targeting ADCs in cases of heterogeneous target expression. We believe our models provide valuable tools for optimizing ADC properties and guiding preclinical and clinical ADC development.

## Acknowledgements

We gratefully thank Anna Kondic for providing feedback on the manuscript.

## Author contributions

NEW: Conceptualization, Model development and implementation, Simulations, Writing – Original draft, Visualization

AC: Simulations, Writing -Review and Editing

MG: Model implementation, Writing -Review and Editing

AVR: Writing -Review and Editing

RS: Model development and implementation

## Funding

This work is funded by Bristol Myers Squibb.

## Competing interests

NEW, MG, AVR are current BMS employees and own BMS stock. RS is a former BMS employee and owns BMS stock. AC was a BMS summer intern.

## Supplemental Figures

**Supplementary Figure 1:**
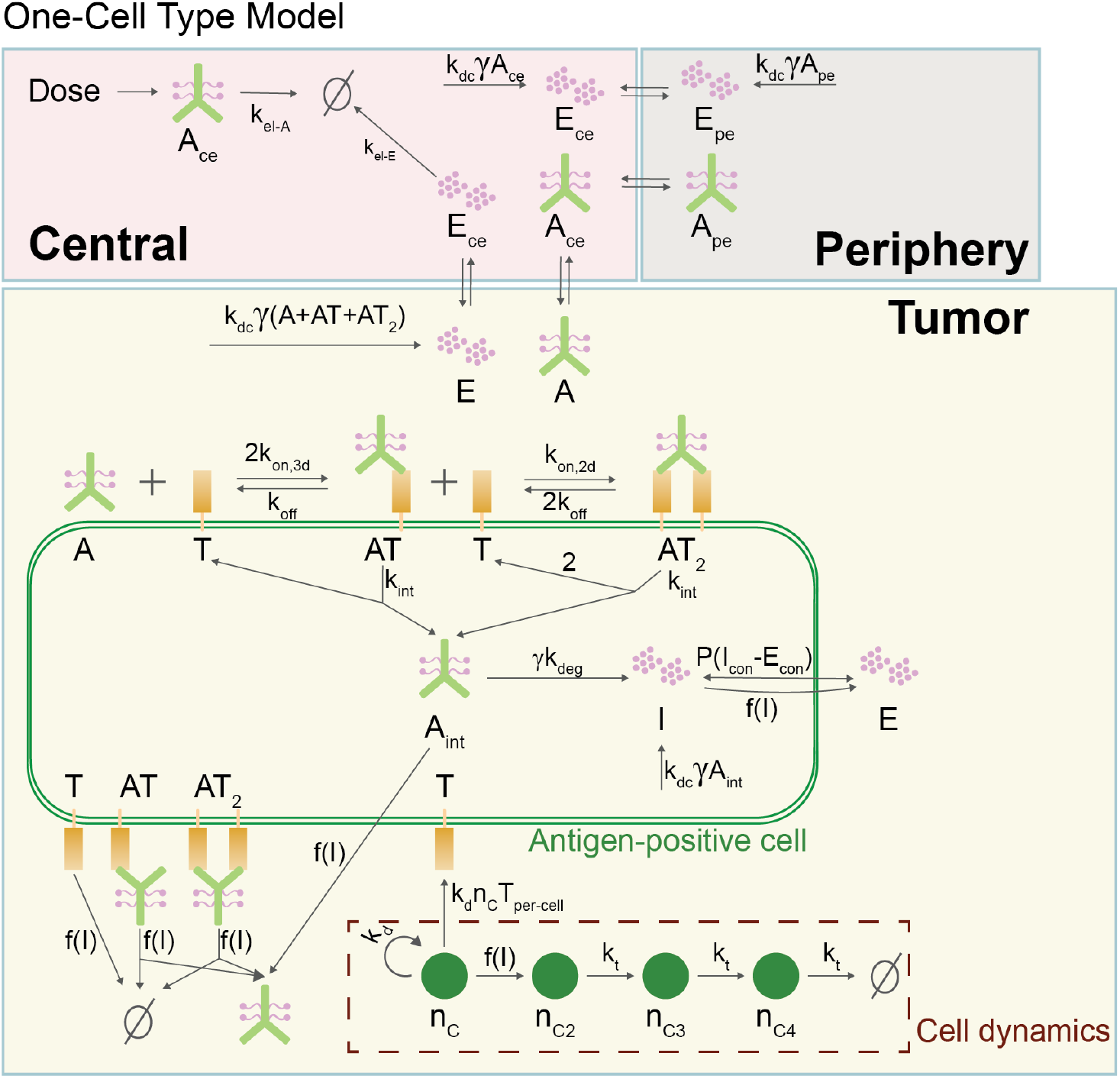
Overview of the One-Cell Type Model. Subscript con denotes concentration. Functions f(I) and f(I_-_) denote sigmoidal functions of the intracellular payload per cell. For details see Supplementary Material.

**Supplementary Figure 2:**
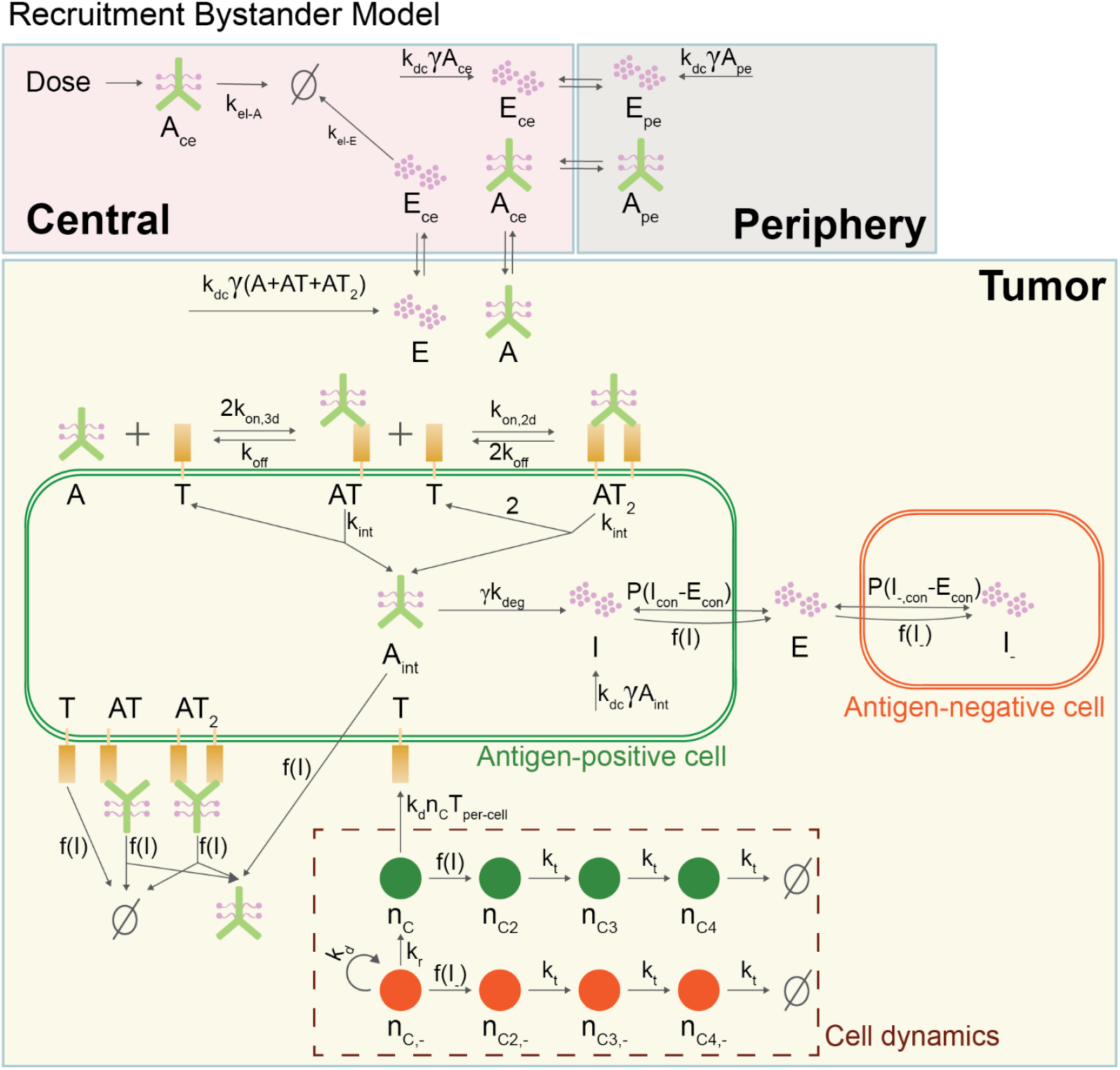
Overview of the Recruitment Bystander Model. Subscript con denotes concentration. Functions f(I) and f(I_-_) denote sigmoidal functions of the intracellular payload per cell. For details see Supplementary Material.

**Supplementary Figure 3:**
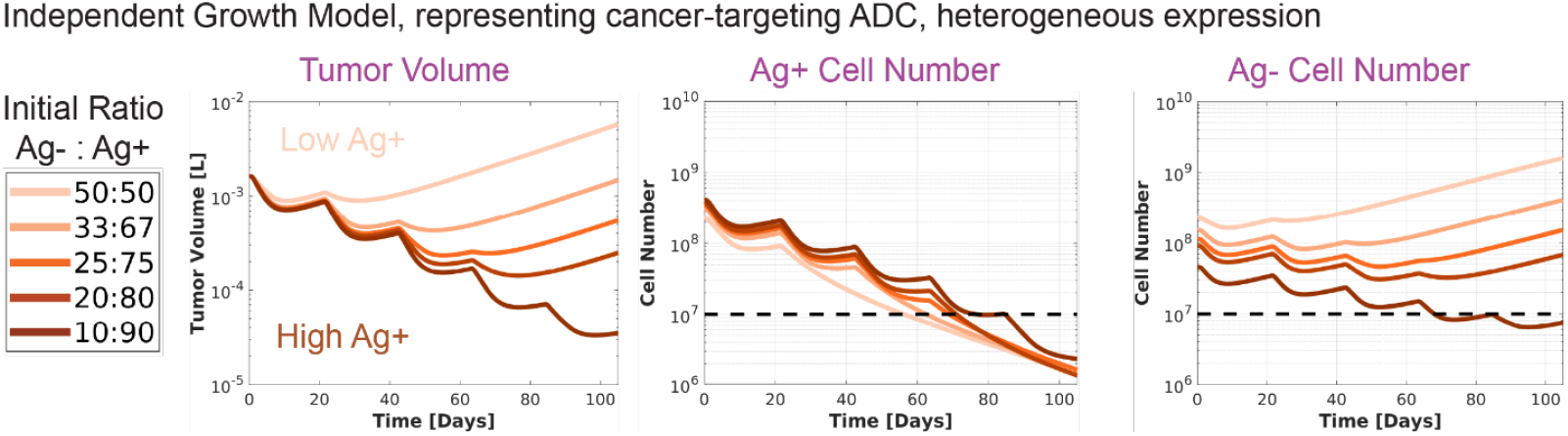
Setting a threshold for Ag-cell number. A threshold is set for Ag-cells, below which the population is assumed unable to recover. This approach can address the limitation of deterministic ODEs in simulating extinction events.

**Supplementary Figure 4:**
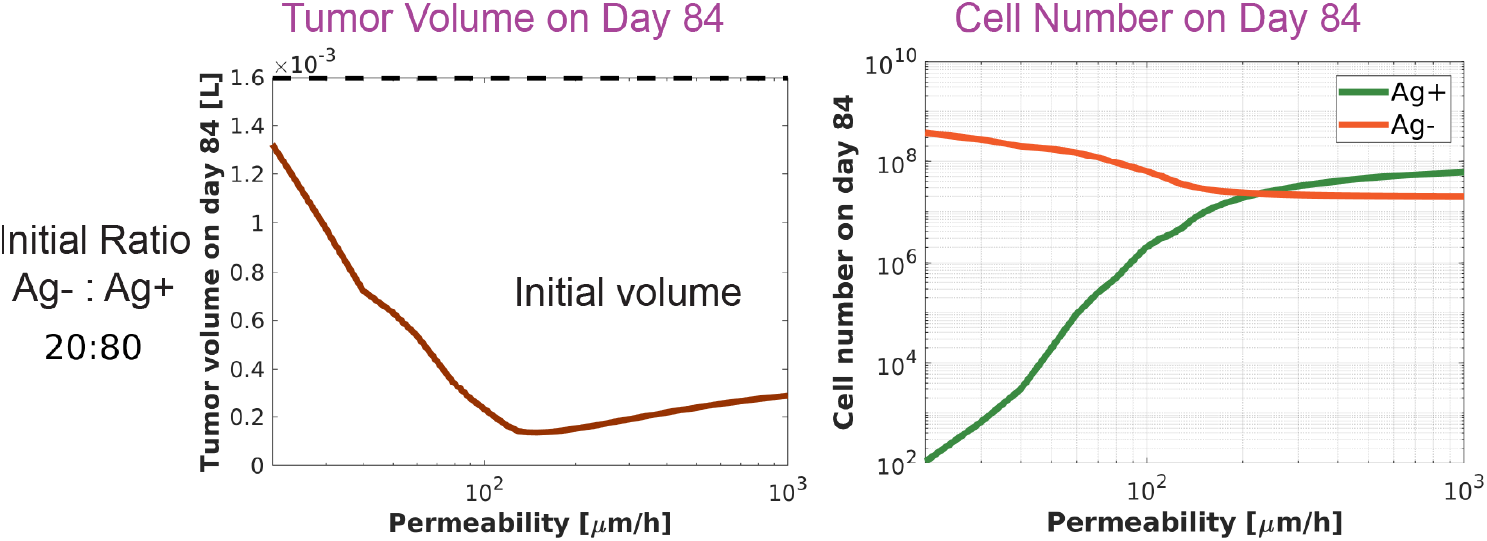
Effect of payload membrane permeability on ADC efficacy. The simulation was run with the Independent Doubling Bystander Model. The parameters are the same as those used in Figure 3, except for initial Ag-to Ag+ ratio. There is an optimal payload permeability that maximizes ADC efficacy on Day 84.

## Supplementary Material

### 1 Models

#### 1.1 State variables

**Table 1:**
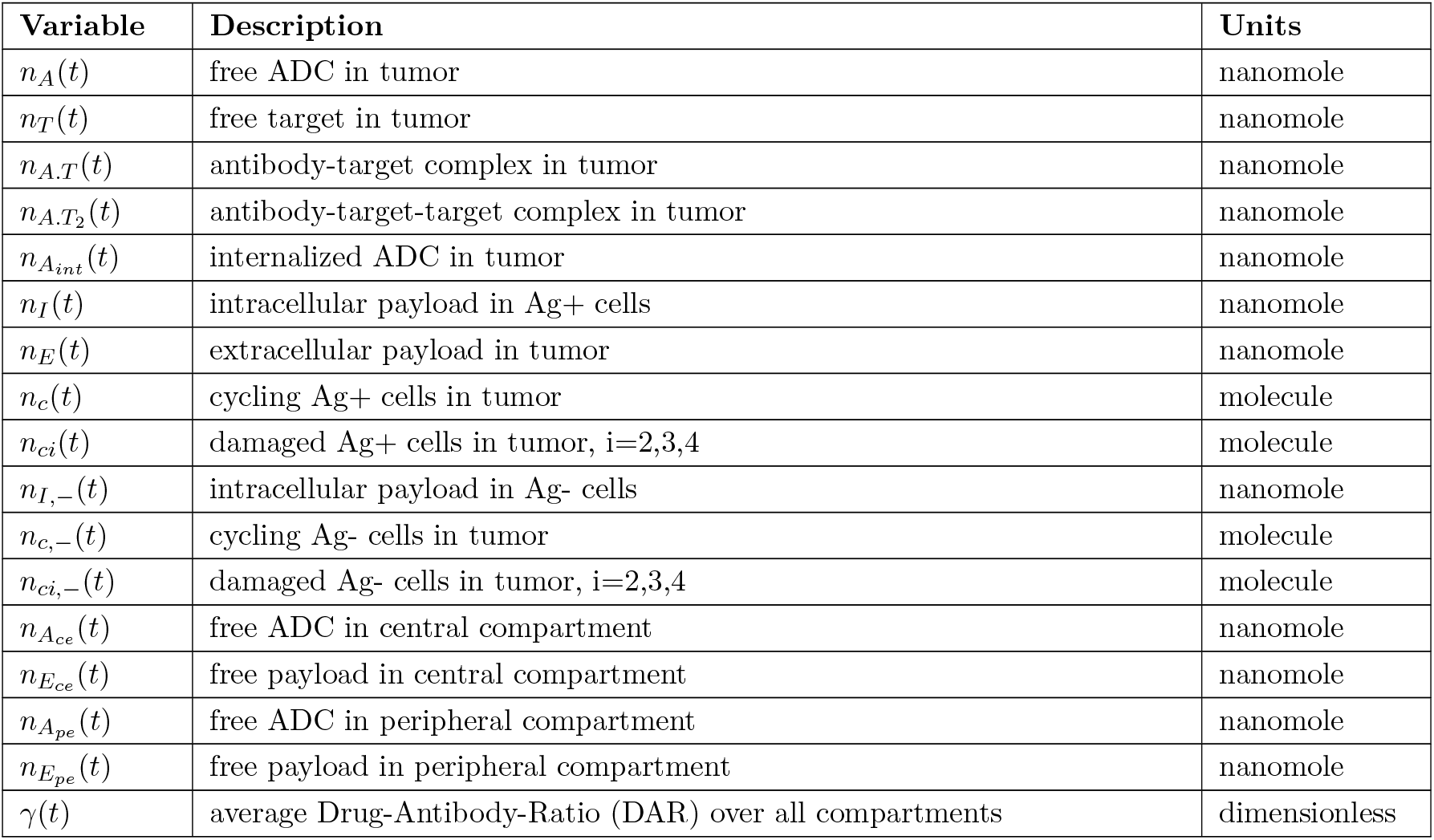
State variables.

| Variable | Description | Units |
| --- | --- | --- |
| $n_A(t)$ | free ADC in tumor | nanomole |
| $n_T(t)$ | free target in tumor | nanomole |
| $n_{A.T}(t)$ | antibody-target complex in tumor | nanomole |
| $n_{A.T_2}(t)$ | antibody-target-target complex in tumor | nanomole |
| $n_{A_{int}}(t)$ | internalized ADC in tumor | nanomole |
| $n_I(t)$ | intracellular payload in Ag+ cells | nanomole |
| $n_E(t)$ | extracellular payload in tumor | nanomole |
| $n_c(t)$ | cycling Ag+ cells in tumor | molecule |
| $n_{ci}(t)$ | damaged Ag+ cells in tumor, i=2,3,4 | molecule |
| $n_{I,-}(t)$ | intracellular payload in Ag- cells | nanomole |
| $n_{c,-}(t)$ | cycling Ag- cells in tumor | molecule |
| $n_{ci,-}(t)$ | damaged Ag- cells in tumor, i=2,3,4 | molecule |
| $n_{A_{ce}}(t)$ | free ADC in central compartment | nanomole |
| $n_{E_{ce}}(t)$ | free payload in central compartment | nanomole |
| $n_{A_{pe}}(t)$ | free ADC in peripheral compartment | nanomole |
| $n_{E_{pe}}(t)$ | free payload in peripheral compartment | nanomole |
| $\gamma(t)$ | average Drug-Antibody-Ratio (DAR) over all compartments | dimensionless |

Ag: Antigen

#### 1.2 Derived variables

**Table 2:** Derived variables.

| Variable | Description | Units |
| --- | --- | --- |
| $n_{c,+T}(t)$ | total Ag+ cells, $n_{c,+T} = n_c + n_{c2} + n_{c3} + n_{c4}$ | molecule |
| $n_{c,-T}(t)$ | total Ag- cells, $n_{c,-T} = n_{c,-} + n_{c2,-} + n_{c3,-} + n_{c4,-}$ | molecule |
| $n_{c,T}(t)$ | total cells, $n_{c,T} = n_{c,+T} + n_{c,-T}$ | molecule |
| $f_{c,+}(t)$ | fraction of Ag+ cells, $f_{c,+} = n_{c,+T}/n_{c,T}$ | dimensionless |
| $f_{c,-}(t)$ | fraction of Ag- cells, $f_{c,-} = n_{c,-T}/n_{c,T}$ | dimensionless |
| $f_{c4,+}(t)$ | fraction of $n_{c4}$ among Ag+ cells, $f_{c4,+} = n_{c4}/n_{c,+T}$ | dimensionless |
| $f_{c4,-}(t)$ | fraction of $n_{c4,-}$ among Ag- cells, $f_{c4,-} = n_{c4,-}/n_{c,-T}$ | dimensionless |
| $V_{tu}(t)$ | tumor volume, $V_{tu} = n_{c,T}V_{cell}/(1 - \epsilon_E)$ | L |
| $n_{I,per-cell}(t)$ | intracellular payload per Ag+ cell, $n_{I,per-cell} = n_I/n_{c,+T}$ | nanomole/molecule |
| $n_{I,-,per-cell}(t)$ | intracellular payload per Ag- cell, $n_{I,-,per-cell} = n_{I,-}/n_{c,-T}$ | nanomole/molecule |
| $V_{tu-eff,A}(t)$ | effective $V_{tu}$ for ADC, $V_{tu-eff,A} = \epsilon_{ADC}V_{tu}$ | L |
| $V_{tu-eff,E}(t)$ | effective $V_{tu}$ for payload, $V_{tu-eff,E} = \epsilon_E V_{tu}$ | L |

#### 1.3 Parameter descriptions

**Table 3:** Parameter Descriptions.

| Parameter | Description |
| --- | --- |
| $MW_{ADC}$ | ADC molecular weight |
| $k_{on,3d}$ | ADC 3-dimensional binding rate (first step) |
| $k_{on,2d}$ | ADC 2-dimensional binding rate (cross-linking) |
| $k_{off}$ | ADC off rate |
| $k_{int}$ | internalization rate |
| $k_{deg}$ | linker cleavage rate + antibody degradation |
| $P_{E,c}$ | permeability of payload across Ag+ cell membrane |
| $P_{E,c-}$ | permeability of payload across Ag- cell membrane |
| $P_S$ | permeability of a species S across tumor capillary |
| $R_{cell}$ | cell radius |
| $S_{cell}$ | cell surface area |
| $V_{cell}$ | cell volume |
| $\rho$ | antibody target number per cell |
| $R_{Krogh}$ | Krogh radius |
| $R_{cap}$ | capillary radius |
| $k_{d,exp}$ | rate of exponential cell growth |
| $k_{d,lin}$ | rate of linear cell growth |
| $\psi$ | parameter controlling the switch between exponential and linear growth |
| $v_{max}$ | maximum killing rate for Ag+ cells |
| $v_{max,-}$ | maximum killing rate for Ag- cells |
| $n$ | hill coefficient for Ag+ cell killing |
| $n_-$ | hill coefficient for Ag- cell killing |
| $IC50$ | payload amount per Ag+ cell for half-maximal killing |
| $IC50_-$ | payload amount per Ag- cell for half-maximal killing |
| $k_t$ | transition rate between cell compartments |
| $\gamma_0$ | starting Drug-Antibody-Ratio (DAR) |
| $k_{dc}$ | deconjugation rate |
| $V_{tu,0}$ | initial tumor volume |
| $\epsilon_S$ | tumor void fraction for species S |
| $V_{ce,S}$ | central volume for species S |
| $V_{pe,S}$ | peripheral volume for species S |
| $CL_S$ | clearance for species S |
| $Q_S$ | intercompartmental clearance for species S |
| $\delta$ | initial ratio of Ag- cells to Ag+ cells for bystander models<br>$\delta$ is assumed to represent the steady state ratio in the absence of treatment. |

S is either ADC or free payload.

#### 1.4 Parameter values

The model is parametrized for Trastuzumab-deruxtecan (T-Dxd). Due to the lack of PK data for deruxtecan (Dxd), the PK parameters for the payload are based on exatecan PK data.

**Table 4:** Parameter Values.

| Parameter | Value | Units | Reference |
| --- | --- | --- | --- |
| $MW_{ADC}$ | 153,702 | g/mol | [1] |
| $k_{on,3d}$ | 2.556 | /nM/h | [2] |
| $k_{on,2d}$ | 731.7 | $\mu\text{m}^2/(\text{molecule}\cdot\text{h})$ | calculated based on [3] |
| $k_{off}$ | 1.26 | /h | [2] |
| $k_{int}$ | 0.1188 | /h | HER2 net internalization [4] |
| $k_{deg}$ | 0.4572 | /h | [1] |
| $P_{E,c}, P_{E,c-}$ | 123.2<br>[20,1000] | $\mu\text{m}/\text{h}$<br>$\mu\text{m}/\text{h}$ | calculated from [4]*<br>varied for Figure 3 |
| $P_A$ | 10.8 | $\mu\text{m}/\text{h}$ | [4] |
| $P_E$ | 3600 | $\mu\text{m}/\text{h}$ | [4] |
| $R_{cell}$ | calculated | $\mu\text{m}$ | $V_{cell} = (4/3)\pi R_{cell}^3$ |
| $S_{cell}$ | calculated | $\mu\text{m}^2/\text{molecule}$ | $S_{cell} = 4\pi R_{cell}^2$ |
| $V_{cell}$ | 2e-12 | L/molecule | [1] |
| $\rho$ | 1e6 | molecule | Her2 per cell [5] |
| $R_{Krogh}$ | 75 | $\mu\text{m}$ | [4] |
| $R_{cap}$ | 8 | $\mu\text{m}$ | [4] |
| $k_{d,exp}$ | 0.0012 | /h | [1] |
| $k_{d,lin}$ | 2.6e-5 | L/h | [1] |
| $\psi$ | 20 | /h | [1], [6] |
| $v_{max}, v_{max,-}$ | 0.01 | /h | T-DXd, N87 cell line [1] |
| $n, n_-$ | 2 | dimensionless | T-DXd, N87 cell line [1] |
| $IC50, IC50_-$ | 2.7e-11 | nanomole/molecule | 13.68nM cellular concentration [1]** |
| $k_t$ | 0.25 | /h | based on 0.17 day delay[1] |
| $\gamma_0$ | 8 | dimensionless | T-DXd [1] |
| $k_{dc}$ | 0<br>[1e-4, 0.05] | /h<br>/h | for all figures except for Figure 4<br>varied for Figure 4 |
| $V_{tu,0}$ | 0.0016 | L | [1]<sup>#</sup> |
| $\epsilon_{ADC}$ | 0.24 | dimensionless | tumor void fraction for ADC [4] |
| $\epsilon_E$ | 0.44 | dimensionless | tumor void fraction for payload [4] |
| $V_{ce,A}$ | 2.77 | L | [7] |
| $V_{pe,A}$ | 5.16 | L | [7] |
| $CL_A$ | 0.0175 | L/h | 0.421 L/day [7] |
| $Q_A$ | 0.0083 | L/h | 0.199 L/day [7] |
| $V_{ce,E}$ | 10.9 | L | exatecan [8] |
| $V_{pe,E}$ | 29.79 | L | exatecan [8] |
| $CL_E$ | 2.78 | L/h | 1.39 L/h/m<sup>2</sup> exatecan [8] |
| $Q_E$ | 7.45 | L/h | exatecan [8] |
| $\delta$ | varied | dimensionless | |

*The payload permeability across cell membrane, *P*_*E*,*c*_, *P*_*E*,*c*−_, is estimated using *k*_*in*_ from [4] as follows:

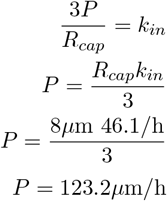

**The payload amount per cell that causes half-maximal killing is estimated using the IC50 concentration reported by [1] as follows:

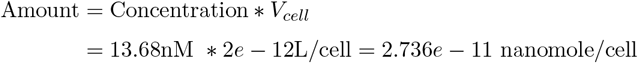

^#^Tumor volume and total cell number are related as such:

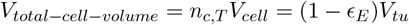

#### 1.5 One-Cell Type Model

##### 1.5.1 Tumor equations

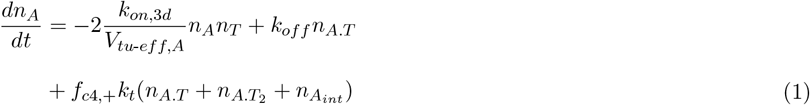

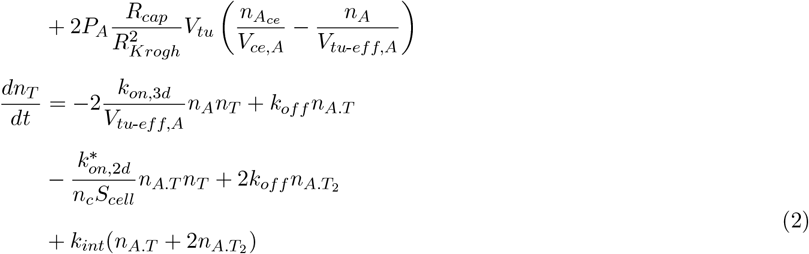

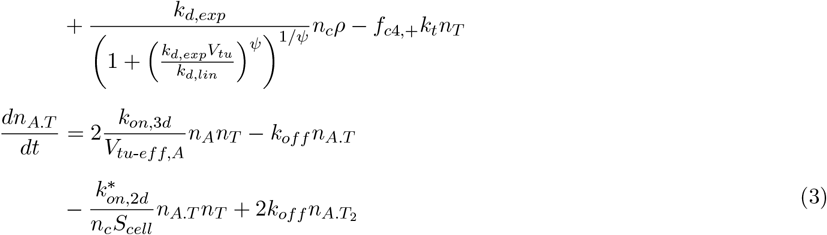

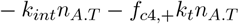

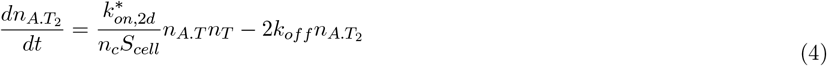

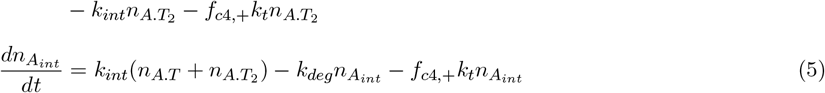

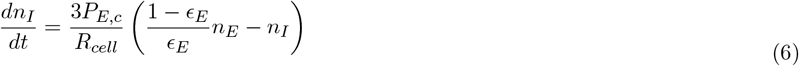

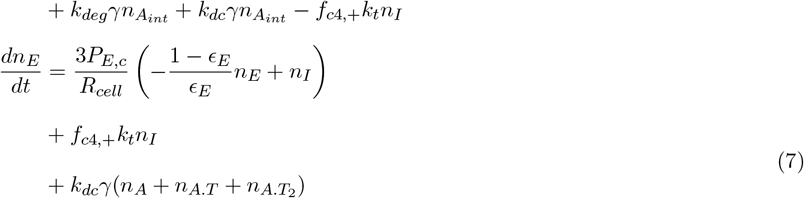

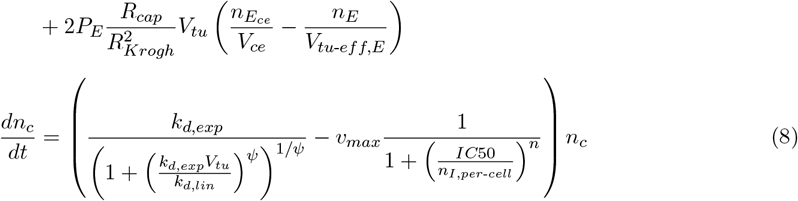

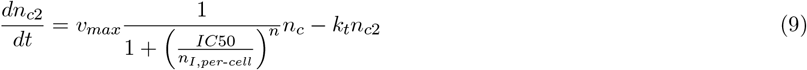

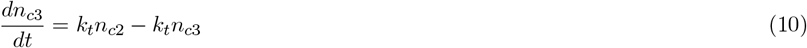

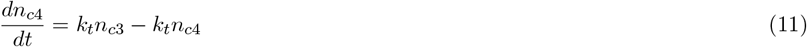

For crosslinking terms, the units of *k*_*on*,2*d*_ is converted from *µ*m^2^/molecule*h to *µ*m^2^/nanomole*h:

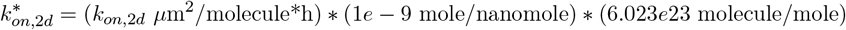

##### 1.5.2 Average DAR equation

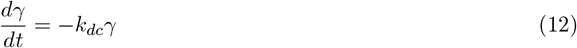

See Section 3 for the derivation of Equation 12.

Note that the average DAR increases with each new dose. Therefore, in multiple dose simulations, the average DAR at the time of dose, *γ*(*t*_*dose*_), is updated using events in SimBiology as follows:

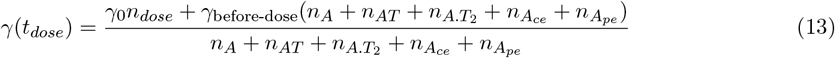

*n*_*dose*_ represents the amount of ADC administered, and *t*_*dose*_ denotes the time of the dosing event.

##### 1.5.3 PK equations

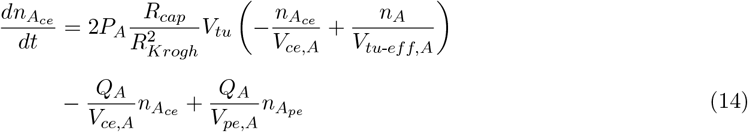

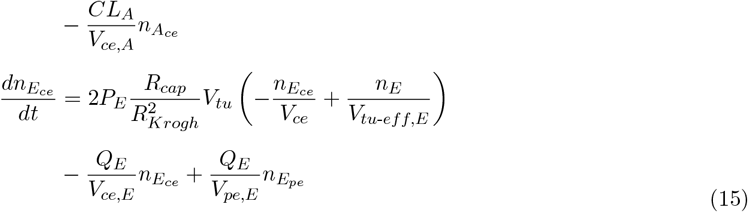

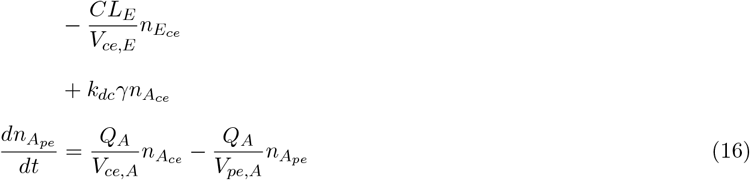

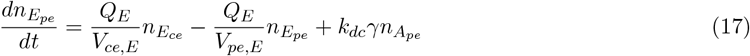

#### 1.6 Extension to bystander models

To explore the bystander effects, we extend the model to include antigen-negative cells. Consequently, equations for intracellular and extracellular payload, i.e. Equations 6 and 7, are modified to include Ag-cells. Additionally, growth equations for Ag-cells are added and the growth equation for Ag+ cells are revised for the Recruitment Bystander Model.

##### Payload equations for bystander models

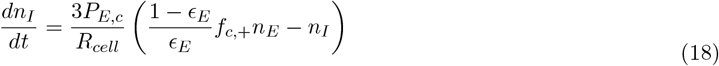

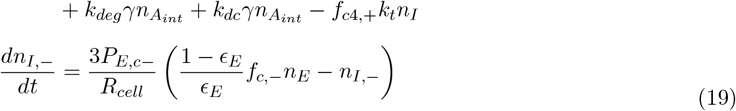

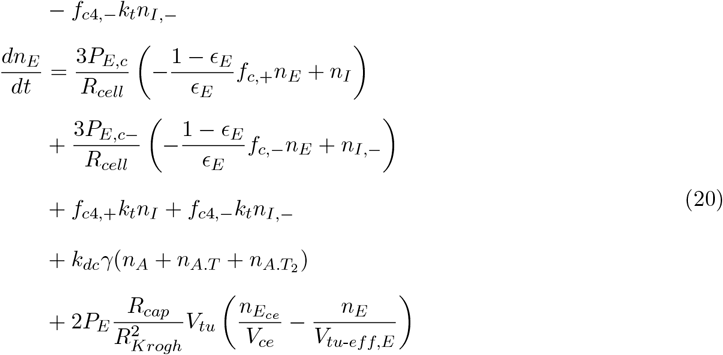

##### Independent Doubling Bystander Model, free target and cell dynamics equations

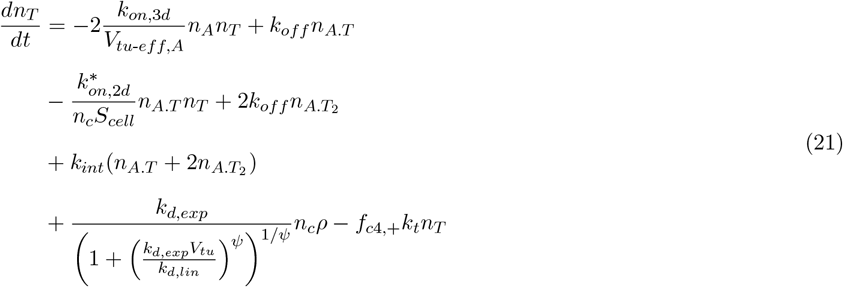

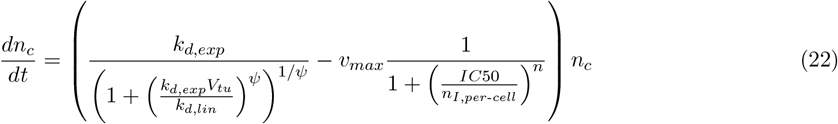

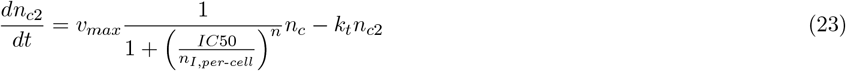

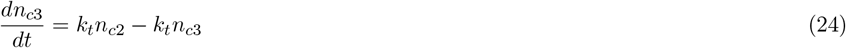

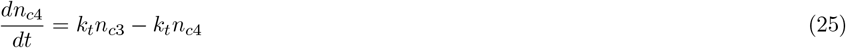

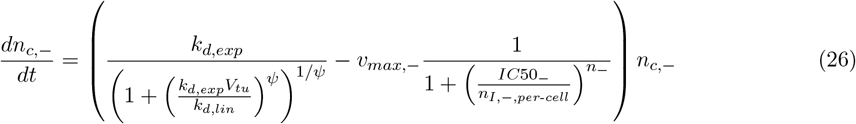

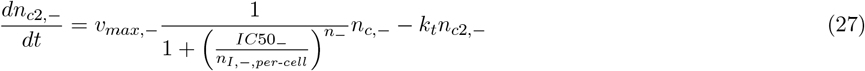

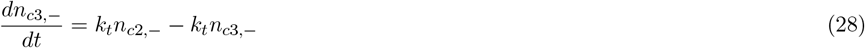

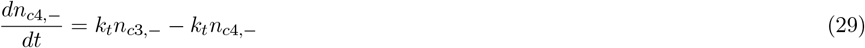

##### Recruitment Bystander Model, free target and cell dynamics equations (see also Section 2)

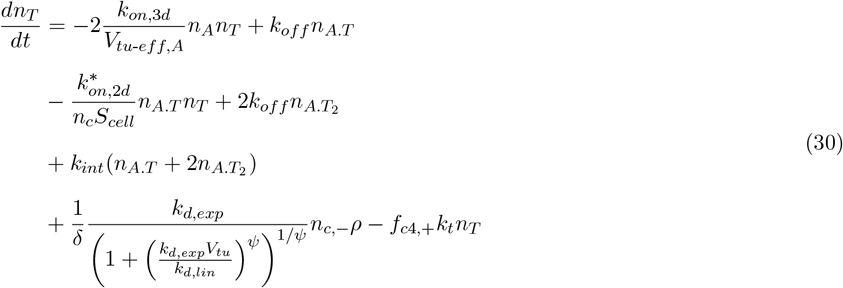

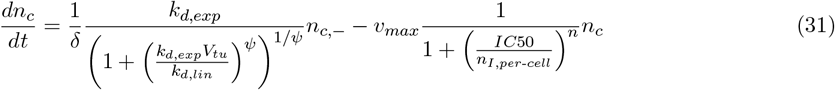

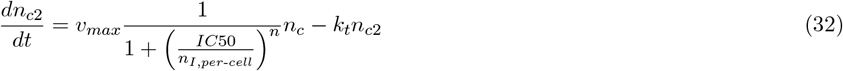

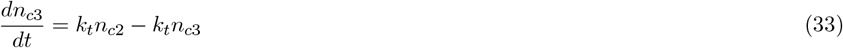

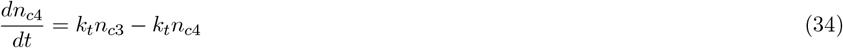

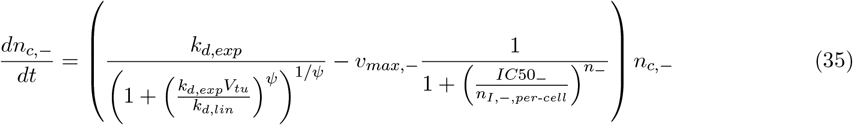

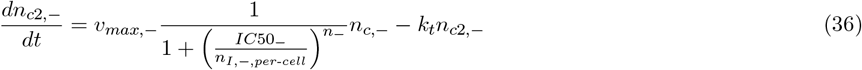

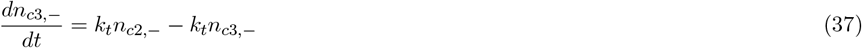

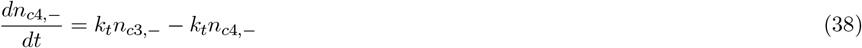

##### Derivation of the terms for payload diffusion across cell membrane

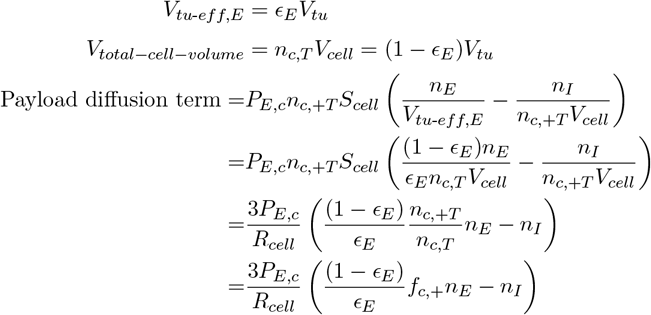

Similarly for antigen-negative cells.

#### 1.7 Initial Conditions

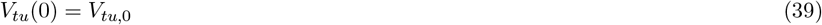

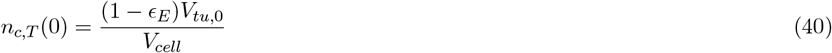

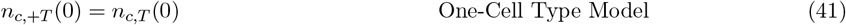

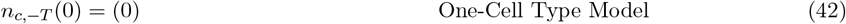

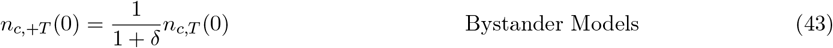

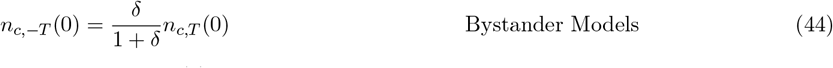

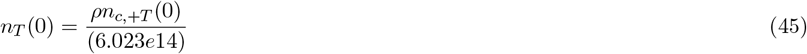

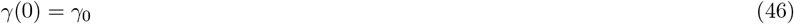

Since *ρ* is in units of molecule/cell, it is divided by the constant 6.023e14 molecule/nanomole. The remaining variables are initialized at a value of 0.

#### 1.8 Model Implementation

The models are implemented in Simbiology and simulated using MATLAB 2022a.

All models are contained within one sbproj model, and three flags control the switch between the models: *flag one cell, flag independent doubling*, and *flag recruitment*. Although *flag independent doubling* is not used in the switches, it is included to make the switching between models more user-friendly: Users can set the flag of the desired model to 1 and the other flags to 0 to switch between models.

The One-Cell Type Model is simulated with *flag one cell* = 1 and *flag recruitment* = 0.

The Independent Growth Bystander Model is simulated with *flag one cell* = 0 and *flag recruitment* = 0. The Recruitment Bystander Model is simulated with *flag one cell* = 0 and *flag recruitment* = 1.

For multiple dose simulations, average DAR increase with each dose, i.e. Equation 13, is implemented using events.

### 2 Modeling antigen-positive cell growth in Recruitment Bystander Model

In this section we motivate the growth term for antigen-positive cells in the Recruitment Bystander Model, as given in Equation 31.

#### 2.1 Exponential growth model

Even though we use exponential-linear growth in the models presented, we first consider exponential growth of Ag-cells, which gives a system that is amenable to an analytical solution, to motivate the form chosen for the Ag+ growth term in the Recruitment Bystander Model.

Let *n*_*c*_ be the number of antigen-positive stromal cells, and let *n*_*c*,−_ be the number of antigen-negative cancer cells. Let *k*_*d*_ be the exponential growth rate for antigen-negative cells, and assume that antigen-negative cells recruit the antigen-positive cells at a rate *k*_*r*_. Additionally, assume that at the initial condition the tumor consists of antigen-negative cancer cells with no antigen-positive stromal cells present. Then, in the absence of treatment, the system is described by the following differential equations:

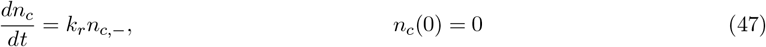

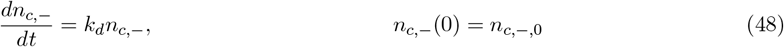

The solutions are given by:

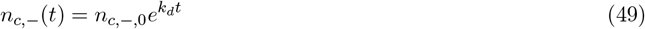

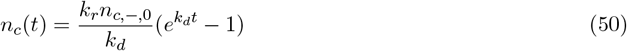

Note that as *t* → ∞, the ratio between antigen-negative and antigen-positive cells approaches a steady-state value determined by the ratio of *k*_*r*_ and *k*_*d*_.

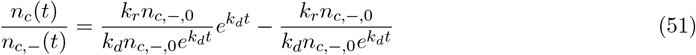

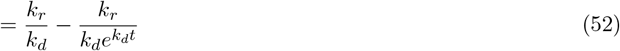

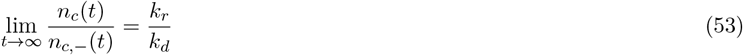

We assume that the initial antigen-negative to antigen-positive ratio, δ, used in simulations are at this steady-state, assuming the tumor has grown for a long time before the start of the treatment. Thus, we set

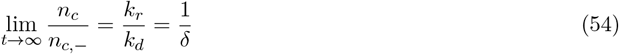

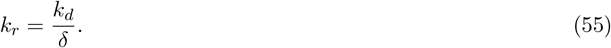

#### 2.2 Exponential/linear growth

Building on the insights from the previous section, we model the growth of cells in the absence of treatment in the Recruitment Bystander Model as follows:

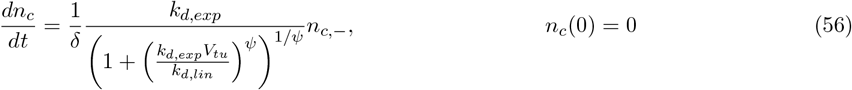

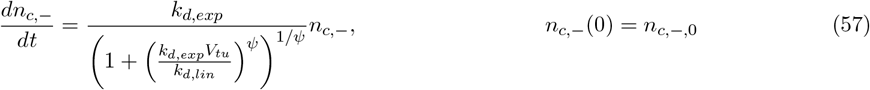

Note that, in this system the ratio, *R* = *n*_*c*_*/n*_*c*,−_, also approaches to 1*/*δ as *t* → ∞:

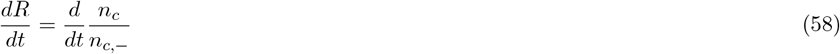

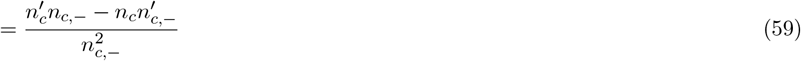

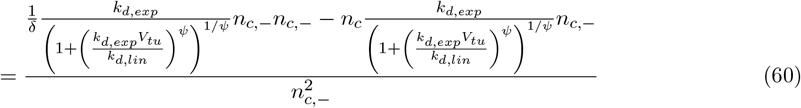

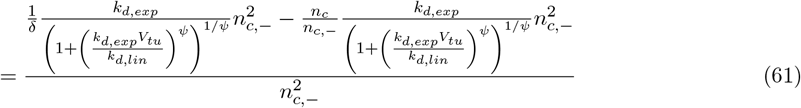

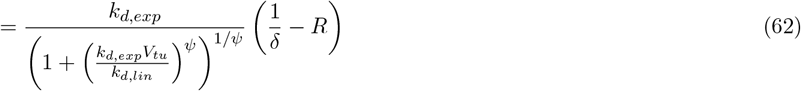

Note that for *R* = 1*/*δ, *dR/dt* = 0. Additionally, for *R <* 1*/*δ, *dR/dt >* 0, and for *R >* 1*/*δ, *dR/dt <* 0. Thus, *R* = 1*/*δ is a stable fixed point.

### 3 Modeling average drug-antibody ratio (DAR)

#### 3.1 Derivation of average DAR equation

Let *n* be the initial DAR of the antibody, and let *A*_*i*_ be antibody with DAR *i*, for *i* = 0, 1, .., *n*. Let *k*_*dc*_ be the rate of deconjugation for one linker-payload. Then,

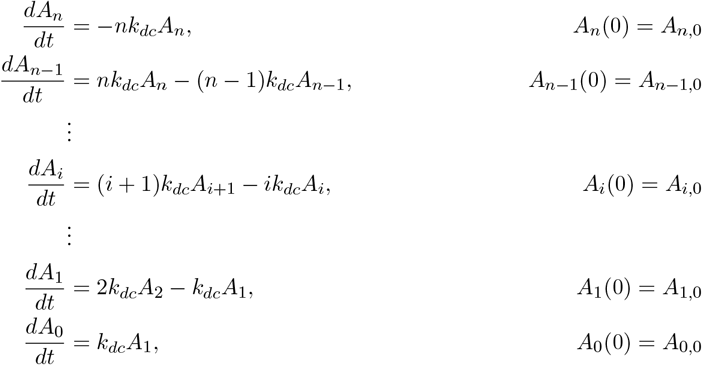

for *i* = 0, 1, .., *n*.

Let *T* be the total antibody, i.e. 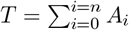.Since 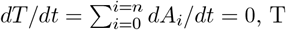 is conserved.

Let *γ*(*t*) be the average drug-antibody ratio (DAR) at a given time:

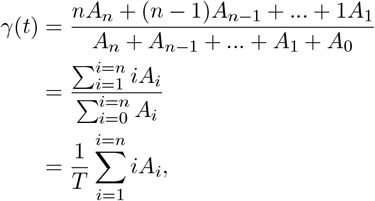

Then,

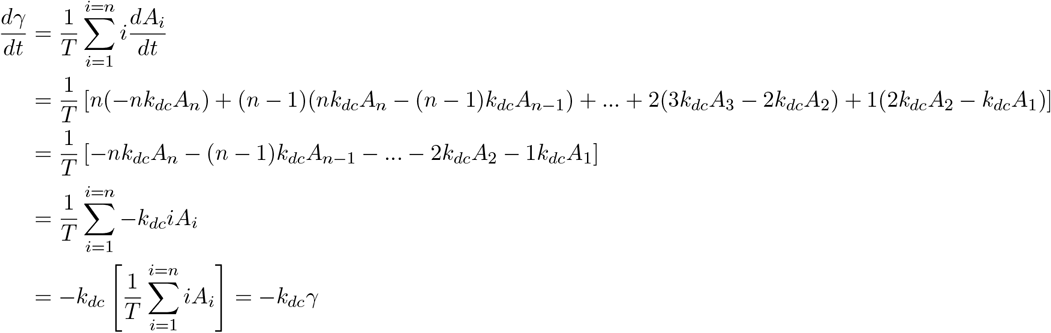

Thus,

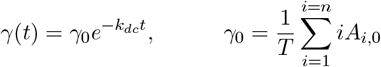

#### 3.2 Released payload

Let *P* be the payload released by deconjugation process. Then,

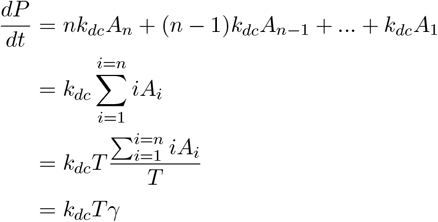

Released payload is incorporated in equations 6, 7, 15, 17, 18, 19, and 20.

#### 3.3 Derivation of average DAR equation with ADC degradation

Note that adding ADC degradation does not change the average DAR equation, assuming that all ADC species of any DAR has the same elimination rate. To show that, let *k*_*el*_ be the elimination rate. Then,

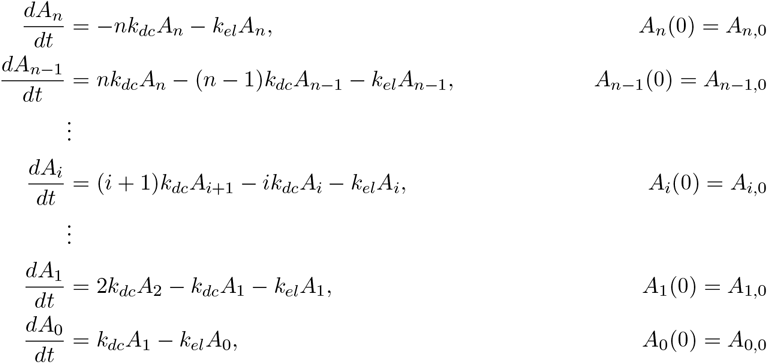

for *i* = 0, 1, .., *n*.

Let *T* be the total antibody. Then,

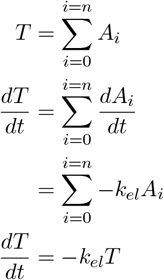

Then average DAR, γ(t), rate of change is given as follows:

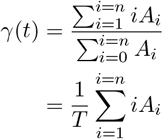

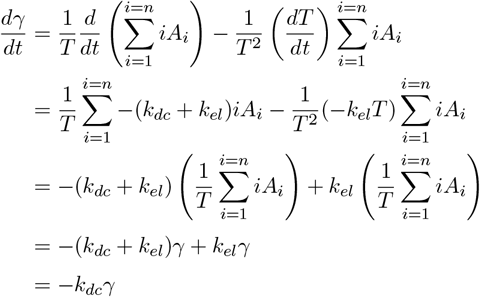

## References

Byun, J. H., & Jung, I. H. (2019). Modeling to capture bystander-killing effect by released payload in target positive tumor cells. BMC Cancer, 19, 1–9.

Colombo, R., & Rich, J. R. (2022). The therapeutic window of antibody drug conjugates: A dogma in need of revision. Cancer Cell, 40(11), 1255–1263.

Colombo, R., Tarantino, P., Rich, J. R., LoRusso, P. M., & de Vries, E. G. E. (2024). The Journey of Antibody– Drug Conjugates: Lessons Learned from 40 Years of Development. Cancer Discovery, 14(11), 2089–2108. 10.1158/2159-8290.CD-24-0708

Crosby, D., Bhatia, S., Brindle, K. M., Coussens, L. M., Dive, C., Emberton, M., Esener, S., Fitzgerald, R. C., Gambhir, S. S., Kuhn, P., Rebbeck, T. R., & Balasubramanian, S. (2022). Early detection of cancer. Science, 375(6586), eaay9040. 10.1126/science.aay9040

Diamantis, N., & Banerji, U. (2016). Antibody-drug conjugates—An emerging class of cancer treatment. British Journal of Cancer, 114(4), 362–367.

Dumontet, C., Reichert, J. M., Senter, P. D., Lambert, J. M., & Beck, A. (2023). Antibody–drug conjugates come of age in oncology. Nature Reviews Drug Discovery, 22(8), 641–661. 10.1038/s41573-023-00709-2

Feng, B., Wu, J., Shen, B., Jiang, F., & Feng, J. (2022). Cancer-associated fibroblasts and resistance to anticancer therapies: Status, mechanisms, and countermeasures. Cancer Cell International, 22(1), 166. 10.1186/s12935-022-02599-7

Frangioni, J. V. (2008). New technologies for human cancer imaging. Journal of Clinical Oncology: Official Journal of the American Society of Clinical Oncology, 26(24), 4012–4021. 10.1200/JCO.2007.14.3065

Fu, Z., Li, S., Han, S., Shi, C., & Zhang, Y. (2022). Antibody drug conjugate: The “biological missile” for targeted cancer therapy. Signal Transduction and Targeted Therapy, 7(1), 1–25. 10.1038/s41392-022-00947-7

Garrison, M. A., Hammond, L. A., Geyer Jr, C. E., Schwartz, G., Tolcher, A. W., Smetzer, L., Figueroa, J. A., Ducharme, M., Coyle, J., Takimoto, C. H., & others. (2003). A phase I and pharmocokinetic study of exatecan mesylate administered as a protracted 21-day infusion in patients with advanced solid malignancies. Clinical Cancer Research, 9(7), 2527–2537.

Grairi, M., & Le Borgne, M. (2024). Antibody–drug conjugates: Prospects for the next generation. Drug Discovery Today, 29(12), 104241. 10.1016/j.drudis.2024.104241

Jordan, M. A., & Wilson, L. (2004). Microtubules as a target for anticancer drugs. Nature Reviews Cancer, 4(4), 253–265. 10.1038/nrc1317

Khera, E., Cilliers, C., Bhatnagar, S., & Thurber, G. M. (2018). Computational transport analysis of antibody-drug conjugate bystander effects and payload tumoral distribution: Implications for therapy. Molecular Systems Design & Engineering, 3(1), 73–88.

Lam, I., Pilla Reddy, V., Ball, K., Arends, R. H., & Mac Gabhann, F. (2022). Development of and insights from systems pharmacology models of antibody-drug conjugates. CPT: Pharmacometrics & Systems Pharmacology, 11(8), 967–990.

Li, F., Emmerton, K. K., Jonas, M., Zhang, X., Miyamoto, J. B., Setter, J. R., Nicholas, N. D., Okeley, N. M., Lyon, R. P., Benjamin, D. R., & others. (2016). Intracellular released payload influences potency and bystander-killing effects of antibody-drug conjugates in preclinical models. Cancer Research, 76(9), 2710–2719.

Nguyen, T. D., Bordeau, B. M., & Balthasar, J. P. (2023). Mechanisms of ADC Toxicity and Strategies to Increase ADC Tolerability. Cancers, 15(3), Article 3. 10.3390/cancers15030713

Purcell, J. W., Tanlimco, S. G., Hickson, J., Fox, M., Sho, M., Durkin, L., Uziel, T., Powers, R., Foster, K., McGonigal, T., Kumar, S., Samayoa, J., Zhang, D., Palma, J. P., Mishra, S., Hollenbaugh, D., Gish, K., Morgan-Lappe, S. E., Hsi, E. D., & Chao, D. T. (2018). LRRC15 Is a Novel Mesenchymal Protein and Stromal Target for Antibody–Drug Conjugates. Cancer Research, 78(14), 4059–4072. 10.1158/0008-5472.CAN-18-0327

Scheuher, B., Ghusinga, K. R., McGirr, K., Nowak, M., Panday, S., Apgar, J., Subramanian, K., & Betts, A. (2022). Towards a platform quantitative systems pharmacology (QSP) model for preclinical to clinical translation of antibody drug conjugates (ADCs).

Shah, D. K., Haddish-Berhane, N., & Betts, A. (2012). Bench to bedside translation of antibody drug conjugates using a multiscale mechanistic PK/PD model: A case study with brentuximab-vedotin. Journal of Pharmacokinetics and Pharmacodynamics, 39, 643–659.

Simeoni, M., Magni, P., Cammia, C., De Nicolao, G., Croci, V., Pesenti, E., Germani, M., Poggesi, I., & Rocchetti, M. (2004). Predictive pharmacokinetic-pharmacodynamic modeling of tumor growth kinetics in xenograft models after administration of anticancer agents. Cancer Research, 64(3), 1094–1101.

Singh, A. P., & Shah, D. K. (2019). A ‘Dual’ Cell-Level Systems PK-PD Model to Characterize the Bystander Effect of ADC. Journal of Pharmaceutical Sciences, 108(7), 2465–2475. 10.1016/j.xphs.2019.01.034

Tomicic, M. T., & Kaina, B. (2013). Topoisomerase degradation, DSB repair, p53 and IAPs in cancer cell resistance to camptothecin-like topoisomerase I inhibitors. Biochimica et Biophysica Acta (BBA) - Reviews on Cancer, 1835(1), 11–27. 10.1016/j.bbcan.2012.09.002

Tsuchikama, K., Anami, Y., Ha, S. Y., & Yamazaki, C. M. (2024). Exploring the next generation of antibody– drug conjugates. Nature Reviews Clinical Oncology, 21(3), 203–223.

van Pelt, G. W., Kjær-Frifeldt, S., van Krieken, J. H. J. M., Al Dieri, R., Morreau, H., Tollenaar, R. A. E. M., Sørensen, F. B., & Mesker, W. E. (2018). Scoring the tumor-stroma ratio in colon cancer: Procedure and recommendations. Virchows Archiv, 473(4), 405–412. 10.1007/s00428-018-2408-z

Xiao, Y., & Yu, D. (2021). Tumor microenvironment as a therapeutic target in cancer. Pharmacology & Therapeutics, 221, 107753. 10.1016/j.pharmthera.2020.107753

Zhang, C., Fei, Y., Wang, H., Hu, S., Liu, C., Hu, R., & Du, Q. (2023). CAFs orchestrates tumor immune microenvironment—A new target in cancer therapy? Frontiers in Pharmacology, 14. 10.3389/fphar.2023.1113378

## References

[1] Bruna Scheuher, Khem Raj Ghusinga, Kimiko McGirr, Maksymilian Nowak, Sheetal Panday, Joshua Apgar, Kalyanasundaram Subramanian, and Alison Betts. Towards a platform quantitative systems pharmacology (qsp) model for preclinical to clinical translation of antibody drug conjugates (adcs). 2022.

[2] Jenny Bostrom, Lauric Haber, Patrick Koenig, Robert F Kelley, and Germaine Fuh. High affinity antigen recognition of the dual specific variants of herceptin is entropy-driven in spite of structural plasticity. PloS one, 6(4):e17887, 2011.

[3] Eric Neil Kaufman and Rakesh K Jain. Effect of bivalent interaction upon apparent antibody affinity: experimental confirmation of theory using fluorescence photobleaching and implications for antibody binding assays. Cancer research, 52(15):4157–4167, 1992.

[4] Eshita Khera, Cornelius Cilliers, Sumit Bhatnagar, and Greg M Thurber. Computational transport analysis of antibody-drug conjugate bystander effects and payload tumoral distribution: implications for therapy. Molecular Systems Design & Engineering, 3(1):73–88, 2018.

[5] Cornelius Cilliers, Hans Guo, Jianshan Liao, Nikolas Christodolu, and Greg M Thurber. Multiscale modeling of antibody-drug conjugates: connecting tissue and cellular distribution to whole animal pharmacokinetics and potential implications for efficacy. The AAPS journal, 18:1117–1130, 2016.

[6] Monica Simeoni, Paolo Magni, Cristiano Cammia, Giuseppe De Nicolao, Valter Croci, Enrico Pesenti, Massimiliano Germani, Italo Poggesi, and Maurizio Rocchetti. Predictive pharmacokineticpharmacodynamic modeling of tumor growth kinetics in xenograft models after administration of anticancer agents. Cancer research, 64(3):1094–1101, 2004.

[7] Ophelia Yin, Yuan Xiong, Seiko Endo, Kazutaka Yoshihara, Tushar Garimella, Malaz AbuTarif, Russ Wada, and Frank LaCreta. Population pharmacokinetics of trastuzumab deruxtecan in patients with her2-positive breast cancer and other solid tumors. Clinical Pharmacology & Therapeutics, 109(5):1314– 1325, 2021.

[8] Mitchell A Garrison, Lisa A Hammond, Charles E Geyer Jr, Garry Schwartz, Anthony W Tolcher, Leslie Smetzer, Jose A Figueroa, Murray Ducharme, John Coyle, Chris H Takimoto, et al. A phase i and pharmocokinetic study of exatecan mesylate administered as a protracted 21-day infusion in patients with advanced solid malignancies. Clinical cancer research, 9(7):2527–2537, 2003.

